# TreePPL: A Universal Probabilistic Programming Language for Phylogenetics

**DOI:** 10.1101/2023.10.10.561673

**Authors:** Viktor Senderov, Jan Kudlicka, Daniel Lundén, Viktor Palmkvist, Mariana P. Braga, Emma Granqvist, Gizem Çaylak, Thimothée Virgoulay, David Broman, Fredrik Ronquist

## Abstract

We present TreePPL, a universal probabilistic programming language (PPL) designed for probabilistic modeling and inference in phylogenetics. In TreePPL, the model is expressed as a computer program, which can generate simulations from the model conditioned on some input data. Specialized inference machinery then uses this program to estimate the posterior probability distribution. The aim is to allow the user to focus on describing the model, and provide the inference machinery for free. The TreePPL modeling language is meant to be familiar to users of R or Python, and utilizes a functional programming style that facilitates the application of generic inference algorithms. The model program can be conveniently compiled and run from a Python or R environment, which can be used for pre-processing, feeding the model with the observed data, controlling and running the inference, and receiving and post-processing the output data. The inference machinery is generated by a compiler framework developed specifically for supporting domain-specific modeling and inference, the Miking CorePPL framework. It currently supports a range of inference strategies—including sequential Monte Carlo, Markov chain Monte Carlo, and combinations thereof—and is based on several recent innovations that are important for efficient PPL inference on phylogenetic models. It also allows advanced users to implement novel inference strategies for models described using TreePPL or other domain-specific modeling languages. We briefly describe the TreePPL modeling language and the Python environment, and give some examples of modeling and inference with TreePPL. The examples illustrate how TreePPL can be used to address a range of common problem types considered in statistical phylogenetics, from diversification and tree inference to complex trait evolution. A few major challenges remain to be addressed before the phylogenetic model space is adequately covered by efficient automatic inference techniques, but several of them are being addressed in ongoing work on TreePPL. We end the paper by discussing how probabilistic programming can facilitate further use of machine learning in addressing important challenges in statistical phylogenetics.

## 1 Introduction

In software for statistical phylogenetics, it has been common practice to implement the entire problem, from model specification to inference machinery, in a general-purpose programming language like C, C++, Julia, or Java. If a larger set of models is supported, then such programs typically allow the user to select the desired model within the predefined space using command line settings—for instance, PhyloBayes [1], RaXML [2] and IQ-Tree [3]. Alternatively, they may allow the user to provide an elaborate configuration file—for instance, MrBayes [4, 5].

In the last decade, advances in statistical phylogenetics software have paved the way for more flexibility and generality. Key developments include the adoption of flexible model specifications in BEAST [6], and the introduction of a model description language grounded in probabilistic graphical models in RevBayes [7, 8]. More recently, we have seen additional phylogenetic modeling languages introduced [9, 10], and a growing interest in leveraging generic statistical modeling and inference frameworks for the analysis of phylogenetic models [11, 12].

### Probabilistic programming languages

Meanwhile, at the intersection of computer science and statistics, a class of tools called *probabilistic programming languages* (PPLs) has matured [13, 14, 15]. A PPL program encodes a probabilistic model of interest using a programming language instead of the language of mathematics. In the context of Monte Carlo-based inference, a PPL program can be thought of as a stochastic program that generates different output on each invocation. If the program is executed an infinite number of times, it defines a probability distribution on the returned values. To make this paradigm even more powerful, we can associate each execution with a likelihood weight, generating a weighted sample from the probability distribution. In Bayesian inference (also called probabilistic inference), we would like the program to specify the posterior probability distribution—that is, the probability distribution of a model conditioned on some observed data. To facilitate this, a PPL typically includes two special probabilistic constructs: (1) `assume` statements (sometimes called `sample`), which describe latent random variables in the model, and (2) `observe` statements, which condition random variables in the model on observed data. These statements also provide convenient checkpoints allowing generic inference machinery, such as sequential Monte Carlo (SMC) and Markov chain Monte Carlo (MCMC) algorithms, to generate and manipulate simulations from the posterior probability distribution. The aim is to allow the user to focus on describing the model, and to leave the estimation of the posterior probability distribution to validated general-purpose inference algorithms. In this way, the user is provided with estimates of model parameters for free. Given a sufficiently powerful programming language, PPLs can express essentially any probabilistic model regardless of the application domain. The modeling flexibility and the potential for automatic inference have inspired PPL work across a wide range of scientific disciplines in recent years. To cite a few applications, probabilistic programming languages have been explored in affective computing (the study of human emotion with computers) [16], in data cleaning [17], and in phylogenetics [18].

### PPLs separate modeling from inference

In the conventional approach to modeling, the initial step involves the derivation of an inference algorithm for a specific model, followed by its implementation using a classical programming language. In the PPL approach, in contrast, the model specification is separated from the implementation of the statistical inference machinery, which is provided automatically by the PPL compiler [19]. This automation opens up the possibility, for example, for the user to try out several different inference strategies for the problem at hand, and to select the best one without modifying the programmatic problem description [20, 21, 22]. Another important benefit of not having to re-implement the inference machinery for every new model is that it reduces the risk of implementation errors. That is, by reusing existing and well-tested code, we can have higher confidence in the result being correct. As a result, we minimize the risk of prolonged arguments over the correctness of the inference for complicated models, as exemplified in statistical phylogenetics by the BAMM debate [23, 24, 25, 26].

While the PPL approach comes with multiple benefits for domain experts, it poses considerable challenges for the developers of the inference machinery. Indeed, it was not until efficient automatic inference techniques were applied to the most powerful PPLs about a decade ago that the interest in using the PPL approach started to pick up significantly [27]. While the last years have seen an explosion in the development of these techniques, and all techniques (if implemented correctly) result in correct inference for all models, it is still far from guaranteed that the inference machinery is efficient for a particular model, especially if the model is complex. Nonetheless, we are now at a stage of development where automatic inference strategies are efficient enough for PPLs to be used productively in addressing many scientific problems, particularly if the model description is structured in a way that is favorable and the inference compiler implements the toolset required to address the specific challenges posed by the model. The coming years will undoubtedly see rapid progress in addressing many of the current constraints, eventually supporting the PPL vision of automatic inference for all models, regardless of how they are described.

### Using PPLs in phylogenetics

Phylogenetic models are complex, and covering the entire model space requires a powerful modeling language. In particular, expressing many common phylogenetic models, such that the entire model becomes available to automatic inference algorithms, requires a language that is universal in the sense that the number of random variables does not have to be fixed a priori. This requirement necessitates stochastic control flow statements, such as stochastic branching, stochastic recursion, or unbounded loops; we will see several concrete examples of this later in the paper. For instance, in a diversification analysis, we are often interested in inferring speciation and extinction rates conditioned on a so-called reconstructed tree, that is, an observed tree of extant (now living) lineages. This incomplete data requires that the automatic inference algorithms can integrate out any unobserved side lineages that died out before the present. A universal PPL expresses such a problem as a simulation over possible extinct side lineages, using stochastic constructs to represent speciation and extinction events in those side lineages, so that the automatic inference machinery can integrate out these simulations [18]. To the best of our knowledge, existing modeling languages designed specifically for phylogenetics are not universal in this sense (see unreleased work on BALI-Phy https://www.bali-phy.org/models.php for a possible exception). Whether or not one wishes to call such languages PPLs is a matter of personal preference; in the following, we will restrict the term PPL to universal languages.

It has become increasingly clear in recent years that automatic and efficient inference for phylogenetic models, especially strategies that scale up to realistic problem sizes, requires a set of special techniques, which are not widely available in existing PPLs. For instance, integrating out random variables (sometimes called Rao-Blackwellization in reference to the Rao-Blackwell Theorem [28]), where possible, is often important for efficient inference. This integration can be done using an approach called delayed sampling [29], which automatically identifies conjugacy relations in the model. Diversification models and similar phylogenetic problems require so-called alignment of checkpoints for inference to be efficient [18]. Strategies and algorithms for automating checkpoint alignment and exploiting it in SMC and MCMC inference were described recently [30]. Another automatic inference technique, which is not yet widely available but can lead to considerable efficiency improvements for some models is the alive particle filter (APF) [31]. APF ensures that automatic SMC algorithms sample more efficiently over models where some outcomes are impossible. Diversification models conditioned on a reconstructed tree represent a good example, as they condition the simulation on side lineages going extinct. Thus, a side simulation surviving to the present has to be discarded, decreasing the efficiency of SMC inference unless the APF is used. Finally, many phylogenetic problems are now conditioned on very large data sets, making it important to leverage parallel computing power where possible [19]. Until now, no PPL has provided a complete set of these improved inference techniques.

### The Miking CorePPL framework

Bayesian statistical phylogenetics typically relies on Monte Carlo algorithms, specifically MCMC (e.g., [32, 5, 1]) or, more recently, SMC (e.g., [31, 33]). In the last few years, optimization or hybrid techniques have also been explored, such as variational combinatorial SMC [34] and variational Bayesian phylogenetic inference [35, 36]. Monte Carlo methods rely on clever and efficient ways of sampling or simulating from the target distribution. In order to implement an inference algorithm in a PPL compiler, facilities must be provided by the compiler for the execution of the probabilistic program to be suspended, likelihood updates to be made, and execution traces (different program runs with random variables and stochastic choices instantiated) to be resampled. To manipulate execution traces, some PPLs—such as WebPPL [21]—rely on transforming the program code to continuation-passing style (CPS) [37, 21, 38], which comes with a performance penalty. In order to minimize the overhead, a selective CPStransformation was investigated in [20]. The results show that selective CPS offers a significant performance gain compared to full CPS on both phylogenetic models, such as the cladogenetic diversification shifts (ClaDS [39]) and constant rate birth-death (CRBD [40, 41]) models, and non-phylogenetic models, such as latent Dirichlet allocation (LDA, [42]). Many of the recent advancements in efficiently compiling probabilistic programs have been developed in an intermediate language called Miking CorePPL [19]. Miking CorePPL is, in turn, implemented within the general compiler framework Miking [43], which is designed for the rapid development of domain-specific language by reuse and composition of language fragments. See https://miking.org/ for details.

A further distinguishing feature of Miking CorePPL is that it has a modular way of expressing Monte Carlo inference algorithms, with most of the compiler being agnostic as to the particular inference engine. Thus, it is easily extensible with new optimizations and inference strategies. Currently supported strategies include importance sampling, the bootstrap particle filter (BPF), the alive particle filter (APF) [31], aligned lightweight MCMC [30, 44], trace MCMC, naive MCMC, and particle MCMC–particle independent Metropolis-Hastings (PMCMC-PIMH, [45]). However, as an intermediate language, Miking CorePPL is not designed to be used by end-users unfamiliar with functional programming and without a deep interest in compiler optimizations. Instead, the intention is that *domain-specific languages* should be designed and built on top of Miking CorePPL, to enable user-friendly environments for specific domain users.

TreePPL is one example of such a domain-specific language, a probabilistic programming language designed specifically for phylogenetics problems and with evolutionary biologists in mind as end users. The TreePPL front-end includes library tools and documentation, which will enable practitioners to write probabilistic programs using the advanced Miking CorePPL inference in the back-end. It focuses on providing maximum expressive power (universality), automatic inference suitable for phylogenetic problems, and a simple syntax that should be familiar to computational biologists. With the publication of this paper, we are inviting the community to try TreePPL.

### Paper structure

The remainder of the paper is structured as follows. In Section 2, we introduce probabilistic programming with some simple examples of how you use it to describe models and inference problems. In Section 3, we give an overview of the TreePPL architecture, and how the program describing the model is used in the TreePPL environment. In Section 4, we use three sets of examples of more advanced models and inference problems to illustrate that TreePPL can already push the envelope in statistical phylogenetics. Finally, in Section 5, we wrap up our presentation with a look towards the future.

## 2 Describing Models Using TreePPL Programs

Probabilistic programming is intrinsically linked to Bayesian modeling, in which the primary objective is to determine the posterior probability distribution *p*(*X* | *Y*) for the latent random variables *X* given observed variables *Y*, applying Bayes’ theorem *p*(*X* | *Y*) = *p*(*Y* | *X*)*p*(*X*)*/p*(*Y*). In this formulation, *p*(*X*) denotes a prior distribution for the latent variables, representing an initial assumption or estimate of these variables’ distribution before incorporating any observations. The term p(Y |X) captures the likelihood of observing Y given specific values of X, and the marginal likelihood *p*(*Y*) = *p*(*Y* |*X*)*p*(*X*)*dX* denotes the model evidence, also known as the normalizing constant.

In probabilistic programming, the model, represented by the joint probability *p*(*X, Y*), is encoded as a computer program. Executing this program performs automatic Bayesian inference, generating either the posterior distribution *p*(*X* | *Y*) for latent random variables or the expected value of a function of interest under this distribution, conditioned on observations of *Y*. To illustrate how this works in practice, consider an example of an unfair coin with an unknown probability of heads, denoted as *p*. Suppose we flip the coin 20 times and record the outcomes, or more precisely, whether each outcome is a head or not. Given these observations, we aim to estimate the probability of heads *p*. Listing 1 demonstrates how to implement this model in TreePPL.

A TreePPL program can have several functions but it needs to have a single function that serves as the entry point, labeled by the keyword `model`. Note that we need to specify the type of the model function’s input variable `outcomes`— a vector of Boolean values denoted by `Bool[]`—and its return type, a real number–––denoted by `Real`. The type specifications come after the name of the variable or the function signature, and are separated from them by a colon. Type specifications are typically needed only when defining functions and their arguments, or when describing user-defined types. In the function body, the `assume` and `observe` statements introduce the latent and observed random variables, respectively, with their distributions specified on the right side of the tilde operator. On line 2, the latent random variable *p* is defined with a prior distribution (uniform on the interval [0, 1]), while lines 3–5 model the sequence of coin flips. The `observe` statement on line 4 clamps observed random variables (outcomes of individual coin flips) to their respective observed values. Our primary interest is the posterior distribution of *p*, which the model function returns on line 6. Any variable or set of variables in the program can be returned, if we are interested in estimating its expectation under the posterior probability distribution. Note that the program does not implement an inference algorithm; rather, it simply encodes the probabilistic model. It does not even specify the observations: the actual data are passed to the model function as an argument, via the parameter named `outcomes`. In this way, we can reuse the same model function for inference problems with different observed data.

**Listing 1:**
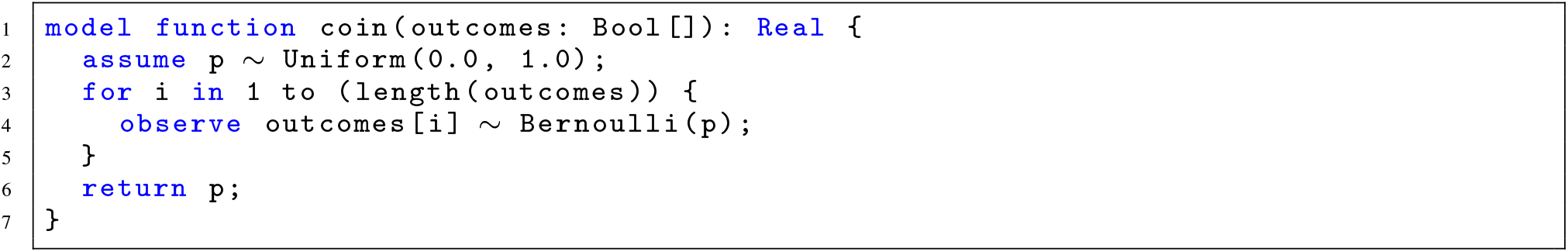
Implementation of coin flipping in TreePPL.

**Listing 2:**
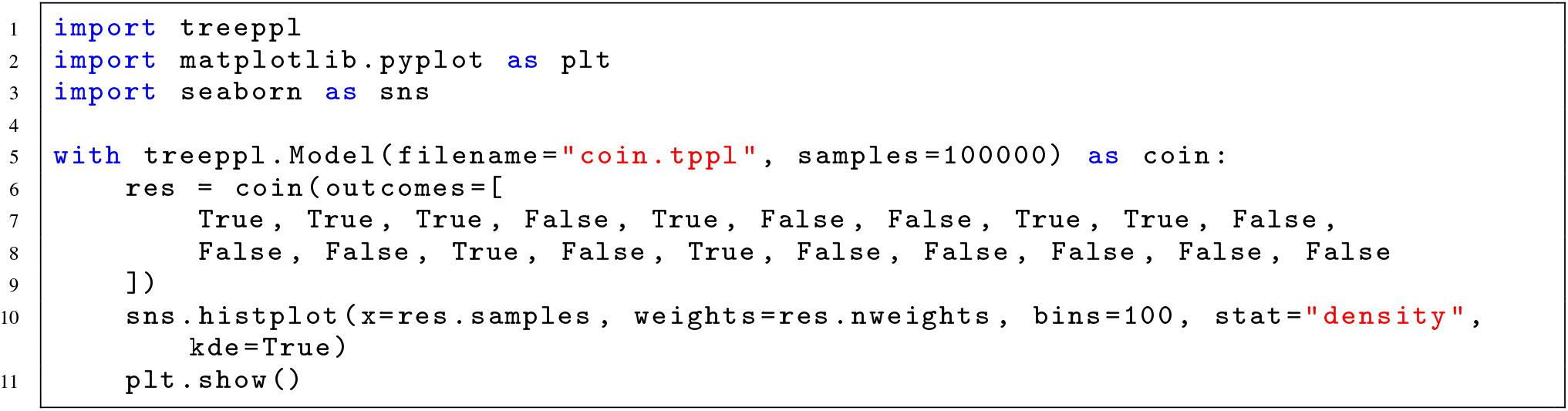
Python wrapper script for executing inference on the coin flipping model and visualizing the posterior distribution.

Although TreePPL programs can be executed directly, a Python interface (and a similar R interface) is available to facilitate data preparation, post-processing, and visualization of inference results (described in more detail in Section 3). Listing 2 demonstrates using Python to perform inference on the coin-flip model and to visualize the posterior distribution.

In this Python program, a `treeppl.Model` object is instantiated on line 5, with parameters indicating the TreePPL model file and inference settings. This model object can be invoked to perform inference (line 6), passing the values for model parameters in TreePPL as named arguments (typically representing the observed data), and returning an inference result object (stored in the variable `res`). This object contains posterior samples, their weights, and an estimate of the model evidence. Line 10 demonstrates plotting the posterior distribution using these samples and weights (Fig. 1).

**Figure 1:**
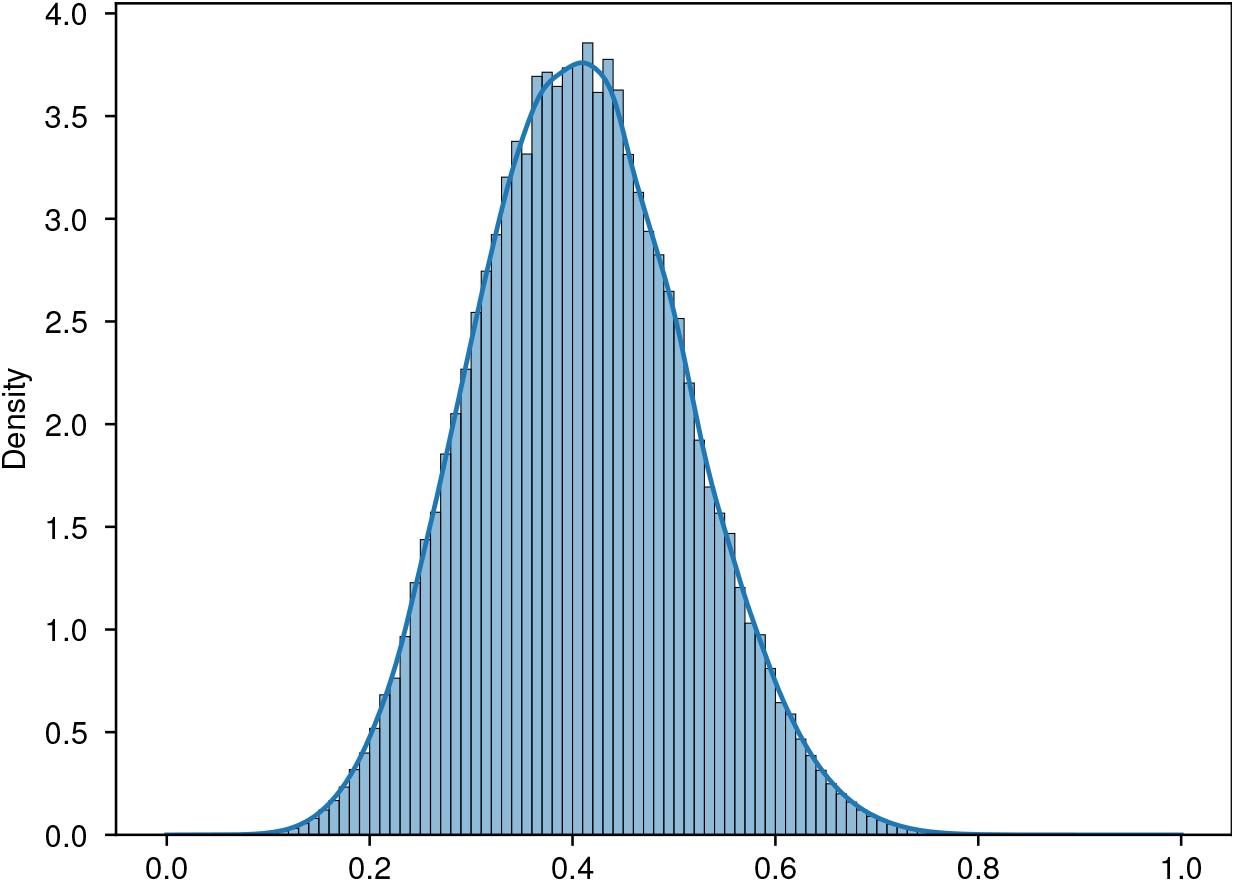
Posterior distribution of the probability of heads for the coin flipping model, with the observed data specified in Listing 2.

Next, let us explore the application of probabilistic programming to phylogenetic modeling. Specifically, let us implement the constant-rate birth-death (CRBD) model. This is an example of a diversification model, an important class of problems focused on inferring the process that generated an evolutionary tree. The CRBD model only has two unknown parameters, the speciation (*λ*) and extinction (*µ*) rates, both of which are assumed to be the same for all lineages and constant throughout evolutionary history. Listing 3 illustrates the generative CRBD model in TreePPL, which simulates a phylogeny beginning with a single species at before-present time `time`, governed by constant speciation rate `lambda` and extinction rate `mu`. The model uses the `Tree` type from the TreePPL standard library. The tree is always in one of the following forms: (1) a `Node`, specifying the right and left descendants (both of type `Tree`), as well as an `age`, a `Real` value specifying the age of the node in user-defined time units; or (2) a `Leaf`, only giving the `age` (0 for extant leaves; a positive value for lineages going extinct before the present). This is an example of the very flexible algebraic data types used in TreePPL (see Section 3.4). On line 2, a waiting time `waitingTime` is sampled from an exponential distribution with rate `lambda + mu`, which is used on line 3 to calculate the event time `eventTime`. If the event time is negative, the species survives to the present time, and the model returns a `Leaf` with `age` 0 (line 5). Otherwise, we determine if the event is speciation (when the boolean variable `isSpeciation`, defined on line 7, is true) or extinction (when `isSpeciation` is false). For a speciation event, the model returns a `Node` with age `eventTime` and recursively generates left and right subtrees, each starting at `eventTime` (line 9). For an extinction event, the model returns a `Leaf` with age `eventTime` (line 15). Recursive capabilities in TreePPL (and other universal probabilistic programming languages) enable the modeling of complex processes such as phylogenetic trees, setting them apart from languages focused on describing probabilistic graphical models (PGMs), which typically are assumed to have a fixed graph structure.

**Listing 3:**
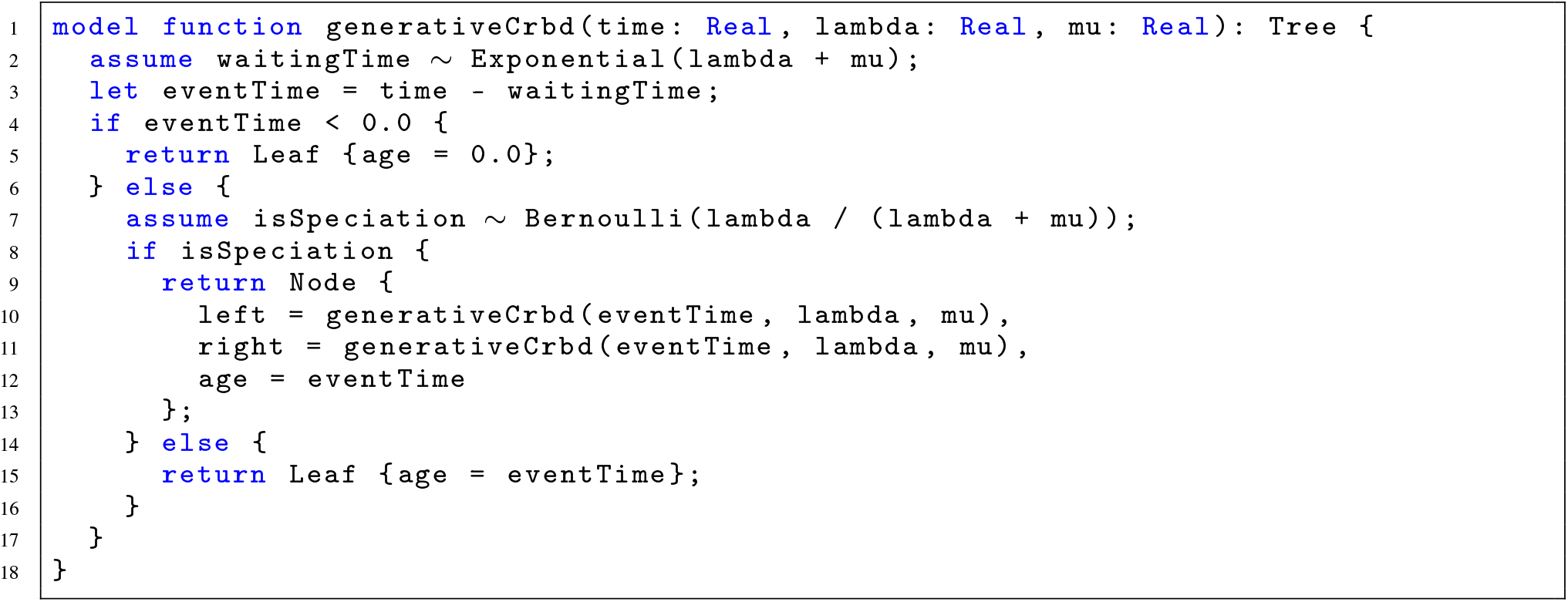
Generative CRBD model in TreePPL.

Finally, let us write a program that allows us to infer the speciation (*λ*) and extinction (*µ*) rates conditional on an observed tree. Typically, the observed tree (“reconstructed tree”) only includes the lineages that survived until the present, that is, we have to accommodate the fact that there might be some unspecified number of unobserved sub-trees (side lineages) that went extinct in the past. In the CRBD model, it is possible to analytically integrate out all possible unobserved sub-trees (see, e.g., the supplementary information in [18]). However, if the speciation or extinction rates are allowed to vary over time or across lineages, as is common in more sophisticated diversification models (e.g., ClaDS [39], BAMM [24], TDBD [46] or LSBDS [47]), analytical solutions are not available, and we instead need to sample over unobserved sub-trees. Here, as a pedagogical example, we will sample over unobserved sub-trees in the CRBD example, even though it is not strictly necessary. Incidentally, by doing so, we also provide a template that can be easily extended to implement a wide range of complex diversification models [18].

**Listing 4:**
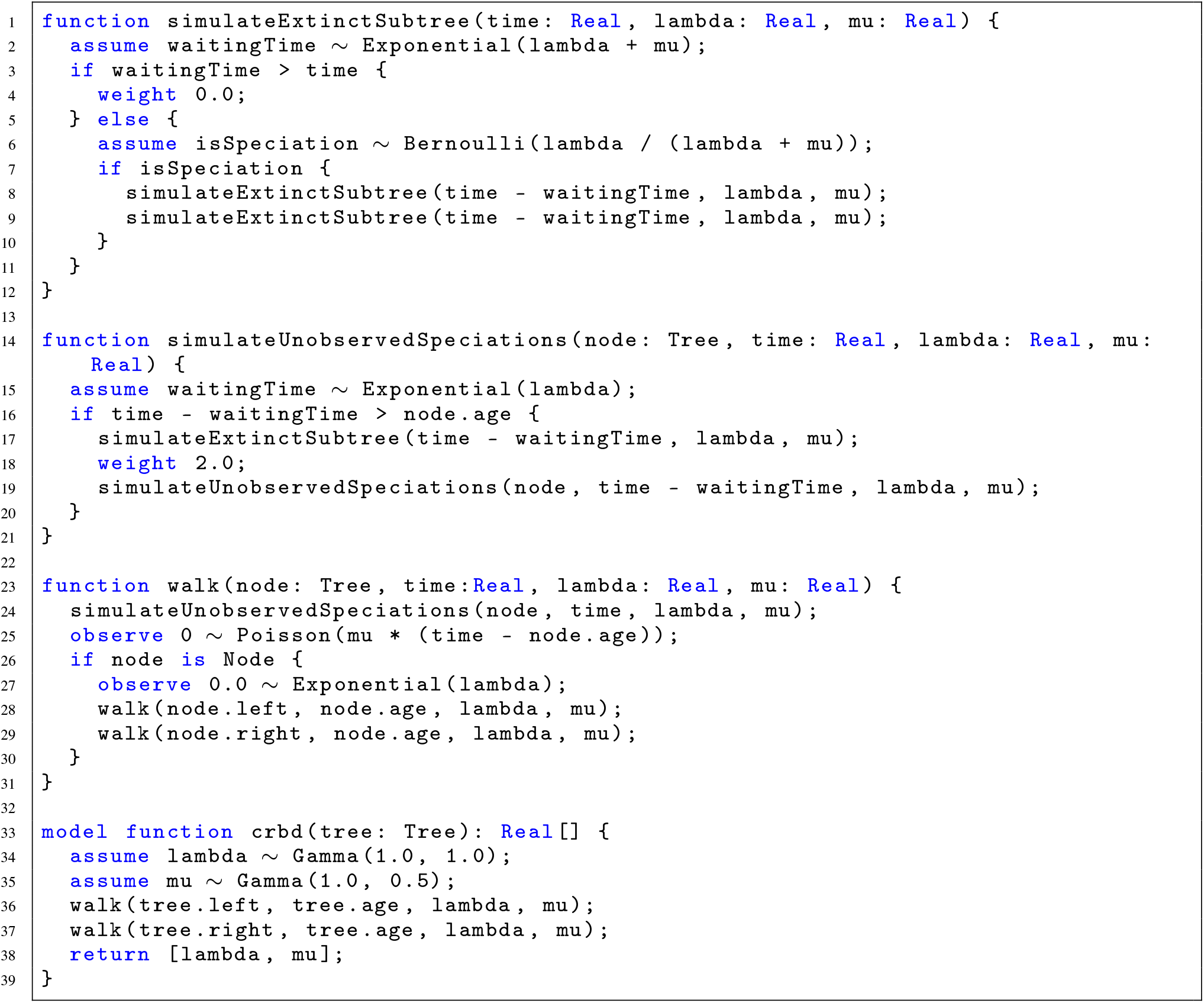
CRBD model implementation in TreePPL.

To specify CRBD inference conditioned on an observed tree, we need to modify and expand the generative model. The complete program contains four functions: the model function, `crbd`, and three helper functions, `walk`, `simulateUnobservedSpeciations` and `simulateExtinctSubtree` (Listing 4). The model function, `crbd`, defines the prior distributions for *λ* and *µ* (lines 34 and 35). It then calls the `walk` function, which recursively traverses the observed tree and simulates the process along its branches, in particular, unobserved speciations (by calling `simulateUnobservedSpeciations`) and extinct subtrees (by calling `simulateExtinctSubtree`) emerging at these speciation events. The Poisson distribution on line 25 represents the number of extinction events along each branch, and is clamped to 0 (if there were any extinction events, the branch would not be a part of the observed tree). The code on line 27 describes the probability of the observed speciation event (in which both descendants survive until the present day) at the end of a descendant branch.

If there is a hidden speciation event on the branch (lines 17–19), the corresponding side lineage is simulated according to the generative model, conditioned on the lineage not surviving until the present, by calling the function `simulateExtinctSubtree` (line 17). For each hidden speciation event, the execution weight is doubled at line 18. This is needed because we do not know whether it was the right or left descendant of the hidden speciation event that survived until the present. This rotational symmetry effectively gives the observed tree two chances of surviving until the present.

The `simulateExtinctSubtree` function is a simplified version of the generative model function in Listing 3 in that it does not store the simulated tree. To enforce extinction of the side lineage, it sets the simulation probability to zero (line 4) if a descendant survives to the present.

This completes the program. To use it for practical inference tasks, it would be run from the Python (or R) environment as described above for the coin flipping model. See below for details (Section 3.2).

For a more extensive discussion of probabilistic programming techniques, we refer to [21, 15], and for a comprehensive TreePPL language reference to our official website https://treeppl.org/docs.

## 3 The TreePPL Programming Environment

In this section, we provide a more in-depth description of the TreePPL architecture, the modeling language and the programming environment. To set up the TreePPL language and environment, please refer to our official website at https://treeppl.org.

### 3.1 Workflow and Architecture

TreePPL consists of the TreePPL compiler and standard library, a Python package, an R package, and a syntax highlighting extension for Visual Studio Code. TreePPL is designed and intended to be used together with Python or R, and we focus on this type of usage in the following. However, we note that it is also possible to run TreePPL as a stand-alone tool, doing the actual TreePPL compilation and execution on the command line. This could be useful in scenarios such as running TreePPL inference on a cluster, where an interactive Python or R environment may not be available.

In Figure 2 we illustrate the software architecture of TreePPL and the recommended development workflow. Using TreePPL for a statistical analysis typically requires the creation of two files by the user: a model file in the TreePPL language (indicated by the “TreePPL program” grey box in Figure 2) and a script in Python or R (indicated by the grey box “Python or R script” in Figure 2). Each model file must contain a model function, whose arguments provide an input interface, and whose return value provides an output interface to the model.

**Figure 2:**
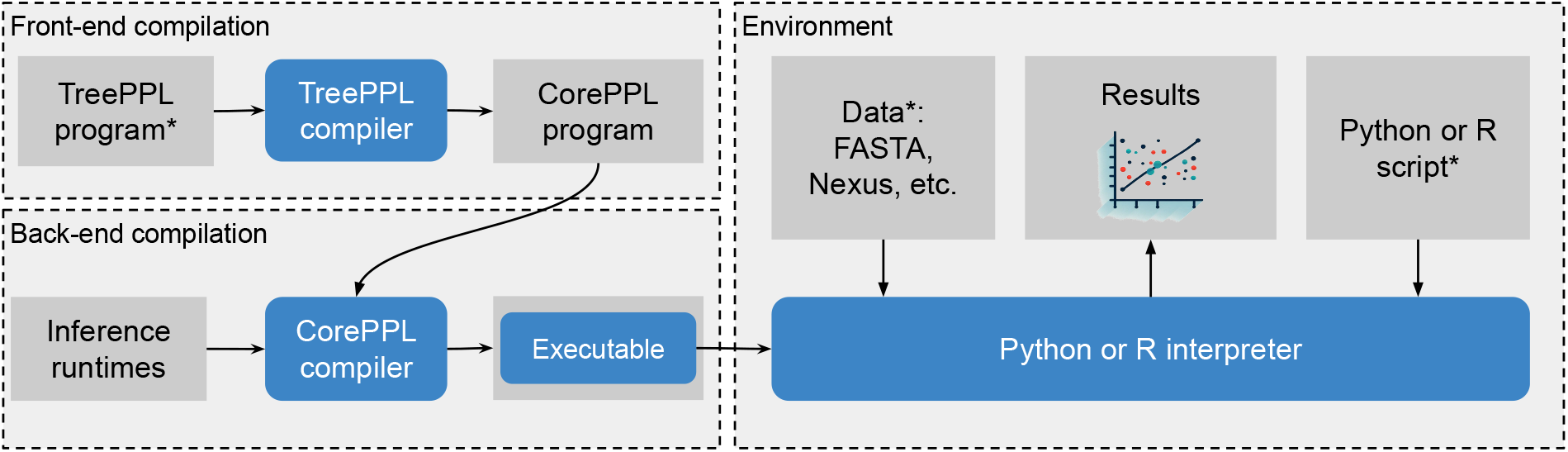
TreePPL programming environment and compiler. The grey boxes represent source code files or data, acting as inputs to or outputs from various workflow stages; the blue boxes represent the compilers or the interpreters involved in acting on the source code and data. With * we have indicated the components that the user supplies, all the others are generated by the programs or supplied with the compilers. The executable is a special case, as it is both the output of the `cppl` compiler and *its output* is processed by the Python or R interpreter to generate plots and do other analysis tasks.

The Python or R script is responsible for:

1. Pre-processing and formatting the input data, such as sequences in FASTA or Nexus format.
2. Selecting one of the available inference algorithms and specifying the inference settings.
3. Running the inference to obtain samples of the posterior distribution of interest.
4. Post-processing the samples and presenting the results.

In the left side of Figure 2, we illustrate the stages through which a TreePPL program goes until an executable is produced. During the initial compilation, the `tpplc` compiler front-end translates the high-level (phylogenetic) model into its intermediate representation, a Miking CorePPL program. The CorePPL compiler, `cppl`, compiles the CorePPL program by applying various optimizations [30, 20]; some of these are applied automatically, while others can be controlled by the user or the Python or R environments. The currently available optimization strategies that can be controlled by the user are:

1. Alignment analysis and placement of resampling points (see checkpoint discussions in Sections 3.3, 4.1), three possibilities:
  a. Where applicable, resample immediately after each checkpoint;
  b. Resample after aligned likelihood updates;
  c. Manual resampling by supplying inference hints;
2. CPS transformation, variants: none, partial, or full (see Section 1);
3. Delayed sampling (in development, see Section 3.3).

The default optimization strategies should work well for most practical analyses; alternative settings are mostly provided for experimental purposes. Please also consult our web page under https://treeppl.org/docs/Reference/ for a complete reference of the various options.

In phylogenetics, as in many other scientific domains, it is not typically possible to sample directly from the posterior distribution of interest. To address this, TreePPL (through the Miking CorePPL back-end) supports a range of inference strategies based on Monte Carlo methods, to perform posterior inference with convergence guarantees. The most basic technique among these is likelihood weighting (a simple form of importance sampling), which draws samples from the prior distribution and weights them according to their posterior probability. This is unlikely to yield a satisfactory sample unless the model is simple and the posterior distribution is similar to the prior distribution.

The more advanced inference strategies can be divided into SMC, MCMC, and mixed SMC/MCMC methods. In SMC algorithms, a set of conditional simulations (often called particles) are run in parallel. During each simulation, the accumulated weight of a particle is computed by summing up log-likelihood values computed at `observe` statements. To improve the accuracy of the posterior estimation, an SMC algorithm also resamples all particles on certain occasions based on their weights (see Section 3.3 for more details). When all conditional simulations have been run to the end, we have a population of particles that represents a valid draw from the posterior probability distribution. The most critical setting in SMC algorithms is typically the number of parallel simulations/particles. The more particles, the better the estimation of the posterior distribution is likely to be.

Among pure SMC strategies, TreePPL currently supports the classical bootstrap particle filter (BPF) as well as the more recent alive particle filter (APF) [31]. The APF is particularly useful for inference problems where particles are likely to ‘die’ because the simulation encounters impossible conditions. For instance, this may occur in a birth-death model, where the simulation of a supposedly extinct side lineage could result in some descendant surviving to the present (see the CRBD example above). The APF can generate better samples of the posterior with fewer particles than BPF when such situations occur frequently. When few or no particles die, the overhead may result in APF being slower than BPF with no gain in sample quality.

MCMC algorithms are the workhorse of statistical phylogenetics, and should be familiar to most empiricists in the field. In PPL implementations of such algorithms, one first obtains a complete sample from the posterior distribution by running a conditional simulation to the end. One then proposes a new sample by modifying the value of one or more of the random variables, and recomputes the posterior probability by re-running the whole or part of the program. Finally, one decides on whether to accept or reject the proposed new sample based on its posterior probability relative to the previous sample, compensating for potential proposal biases (Metropolis-Hastings algorithm). The algorithm generates a Markov chain of samples converging onto the posterior distribution; the quality of the inference is related to the rate of convergence and the length of the Markov chain.

TreePPL supports MCMC algorithms that propose new values of random variables by redrawing from the prior (the distribution associated with each random variable), or by applying a drift kernel to the current value. Two supported approaches are ‘naive MCMC’, which recomputes the entire downstream simulation after modifying a random variable, and ‘trace MCMC’, in which the new proposed sample is a complete rerun of the entire simulation. Naive and trace MCMC are provided mainly for experimental purposes, even though they may be effective for simple real-world inference problems. TreePPL also supports a more advanced MCMC method, called *aligned lightweight MCMC*. See Section 3.3 for more details.

Finally, TreePPL currently provides one algorithm mixing MCMC and SMC strategies, namely particle MCMC–particle independent Metropolis-Hastings (PMCMC-PIMH, [45]). This is perhaps the simplest version of an SMC-within-MCMC approach, where each proposal in the MCMC is a full set of samples (particles) generated by SMC. Future work will add more sophisticated versions of SMC-within-MCMC strategies, as well as MCMC-within-SMC strategies like particle rejuvenation. We also plan to add strategies utilizing tempering [48] (change of the ‘temperature’ of distributions) to improve the quality of the sample. These techniques include, among others, Metropolis-coupled MCMC [49]—familiar to phylogeneticists through its early adoption in MrBayes [50]—and Sequential Change of Measure (also known as ‘SMC sampler’ algorithms) [51, 52]. For an up-to-date account of available TreePPL inference strategies, as well as advice on how to assess the quality of the sample and choose the best strategy for a particular inference problem, please visit https://treeppl.org/docs/Reference/inference.

**Listing 5:**
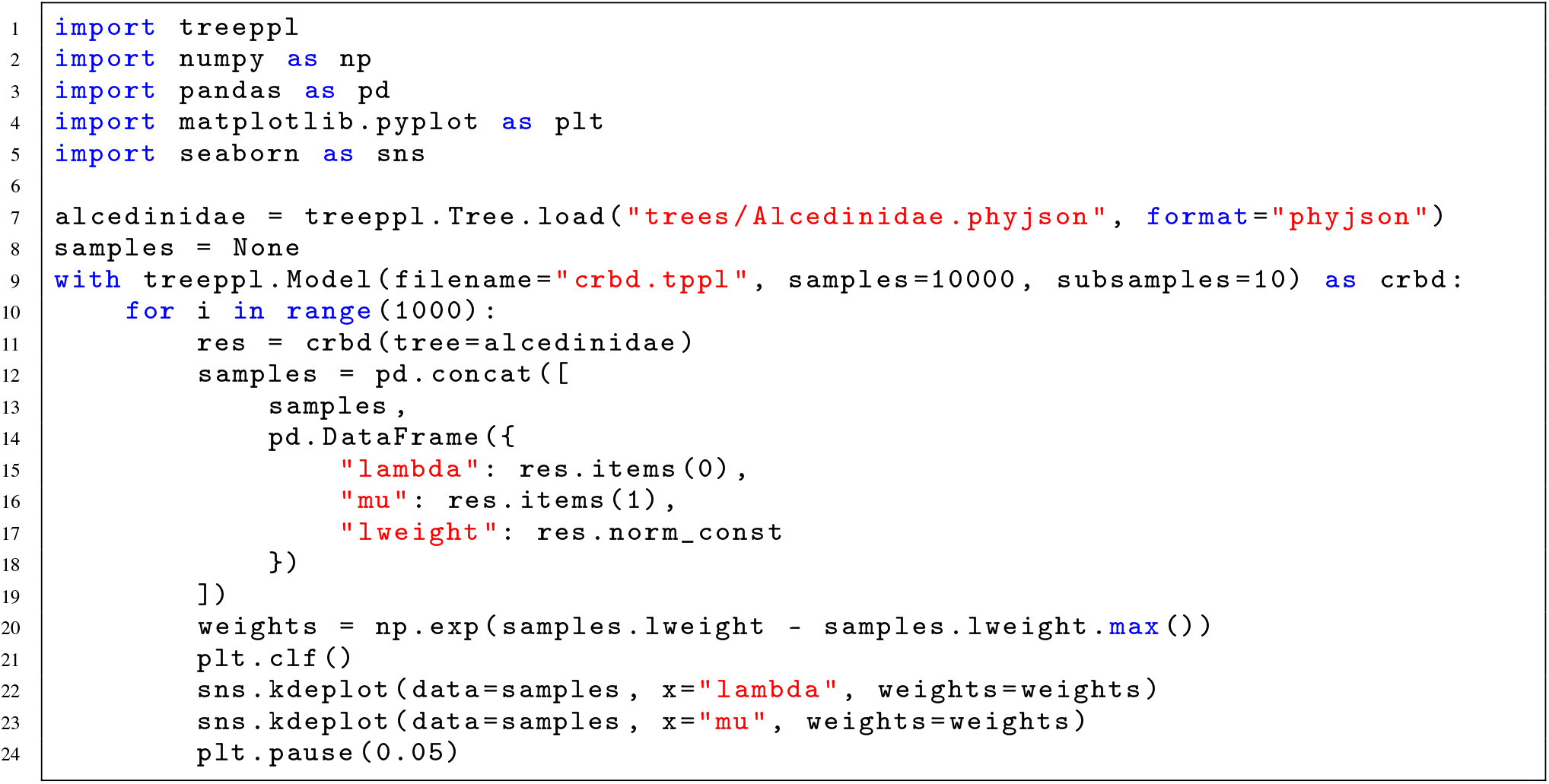
CRBD model implementation in TreePPL/Python (Python part)

### 3.2 Pre- and post-processing with Python or R

The Python package^3^ that comes with TreePPL serves as a versatile interface for creating and executing inference on TreePPL programs. It offers users the advantage of working with input data and processing inference results using a familiar language that comes with a wide range of pre- and post-processing capabilities. These include popular libraries, such as `Biopython`, `Numpy`, `Pandas`, `Matplotlib` and `Seaborn`. An R package with similar functionality is also available^4^.

Listing 5 exemplifies the use of the Python interface for leveraging the CRBD model implemented in TreePPL (Listing 4) to estimate the posterior distribution of the speciation rate.

The Python program first reads in the observed tree data from a PhyJSON^5^ file, a TreePPL-specific file format (line 7). All common data and tree formats available through `Biopython`, such as Nexus and Newick, are also available to TreePPL users.

Next, a `treeppl.Model` instance is created (line 9). This step involves specifying the name of the TreePPL model file (`filename`), the number of samples to be drawn (`samples`) and the number of subsamples to be returned during the inference (`subsamples`). The instantiation process includes reading and compiling the TreePPL model. Note that various other parameters are supported, including specifying the source code as an inline string (`source`), alternative inference methods (`method`; `smc-bpf` by default) and additional parameters for the `tpplc` compiler.

The model instance can be called directly (line 11) to perform inference on the TreePPL model, passing the input data as named arguments that align with the parameters of the TreePPL model function. The Python package encodes the provided input data into the JSON format, which is then passed to the compiled model executable. Subsequently, the output of the model executable, also in JSON format, is parsed. The resulting output includes samples, weights, and estimates of the marginal likelihood (for relevant inference methods), which are returned as a `treeppl.InferenceResult` instance.

In this example program, the SMC-based inference is repeated 1 000 times (line 10). In each iteration, the returned subsamples are added to a Pandas data frame (lines 12–19), and used to update the estimate of the posterior distribution shown to the user (lines 22–23). By repeating the inference many times, as in this example, we increase the accuracy of the inference result.

As shown in this example, Python visualization libraries can be used to plot the resulting posterior distributions. In Figure 3, we have repeated the analysis twice with two different inference methods—the bootstrap particle filter [53] and the alive particle filter [54].

**Figure 3:**
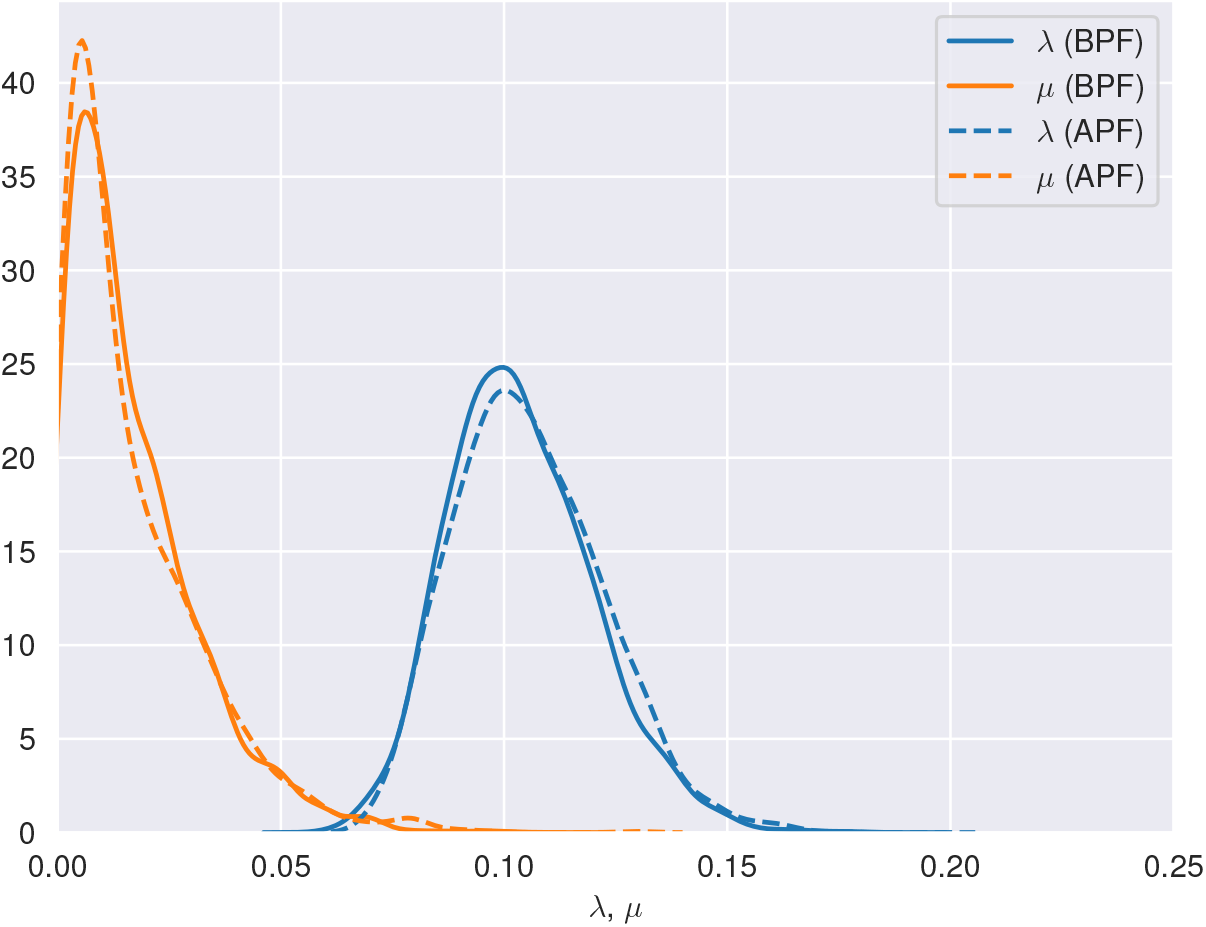
Posterior distributions of the speciation rate *λ* and extinction rate *µ* under the CRBD model inferred using two different inference methods (`smc-bpf` and `smc-apf`).

### 3.3 Optimizations and Inference

In order to make the inference more efficient, the compiler will attempt to optimize the code based on the settings provided by the user (see Section 3.1). However, the model program itself can be structured by the user in ways that may result in important gains in the efficiency of the automatically generated inference machinery, particularly given the constraints of the techniques that are currently available.

#### Automatic alignment and selective CPS

A central concept when compiling PPL programs is a *checkpoint*. At checkpoints, simulations can stop, make modifications, and then decide whether to continue. For instance, in SMC, the essential checkpoint is at `observe` statements, where decisions are made whether to resample or not. In contrast, for MCMC algorithms, the essential checkpoint is at `assume` statements. There are several approaches for handling checkpoints in universal PPLs, e.g., using specific control-flow graphs [19], or to use continuation-passing style (CPS).

TreePPL makes use of many of the state-of-the-art optimization methods recently implemented in the Miking CorePPL framework. One such method is selective CPS [20], which avoids CPS overhead by determining—using static analysis—which parts of a program that need suspension at certain checkpoints.

Another optimization technique is automatic alignment [30]. This approach can optimize both SMC and MCMC algorithms, but in different ways. For SMC, it automatically decides where to perform resampling (at aligned `observe` or `weight` statements, that is, statements that are guaranteed to be executed in the same order, by all simulations). In the case of MCMC, it is used to synchronize and reuse aligned random draws between simulations. In contrast to previous approaches, this aligned lightweight MCMC method [30] avoids the need and overhead of a dynamic database, something that was required in the original lightweight MCMC approach [44].

#### Delayed sampling and belief propagation

In the PPL literature, `assume` statements are also called `sample` statements because they often involve direct sampling of the random variable. However, in many cases it is advantageous not to sample the value of the random variable directly, justifying the syntax adopted by TreePPL. Instead, one can either represent the possible values as a probability vector over outcomes (for numerical integration or summation), or defer sampling until it is possible to determine whether any conjugacy relations allow the variable to be integrated out analytically. The former is closely related to belief propagation, an example of which is Felsenstein’s pruning algorithm [55], frequently used in statistical phylogenetics. The latter is a more recent technique in PPL inference algorithms, referred to as delayed sampling [29], although the exploration of conjugacy relations is as old as Bayesian inference itself. Both methods are known to dramatically accelerate the performance of inference algorithms in many cases. Conjugacy-utilizing optimizations are ideally done by the compiler, and the end-user does not have to worry whether the generative model described with `assume` actually samples the random variables, or instead uses a different technique to achieve the desired outcome. Automatic application of delayed sampling [56] and Felsenstein’s pruning algorithm [57] are both work in progress in TreePPL.

#### Extending the inference

Inference can be controlled by selecting an inference strategy and changing algorithm settings using the Python or R environment, as illustrated above. If the user wants to implement a novel inference strategy, this can be done by writing inference code at the Miking CorePPL level. In order to do this, the user must implement both a runtime library (cf. Figure 2) and a compiler routine for the inference strategy [20]. There are a number of reusable Miking CorePPL compiler and runtime components, which the user may take advantage of in their inference strategy implementation. Some examples of such components are continuation-passing style transformations and automatic alignment. The user may also use existing inference strategy implementations as templates. Please see https://miking.org/ for details.

### 3.4 TreePPL Language Features

Many TreePPL language features should be familiar to users with some previous programming experience. For instance, the syntax follows a brace-bracket style in the tradition of C, C++, etc. We can also comment in the usual C-style with // or /* … */. We refer the reader to the TreePPL website, https://treeppl.org/docs for a detailed language reference, and a set of examples of both general modeling and phylogenetic modeling in TreePPL. Here, we just highlight a few powerful language features, which may be unfamiliar to potential users but that facilitate the description of phylogenetic models and the application of automatic inference.

#### Functional programming and immutability

The TreePPL language is a functional programming language. From a practical coding perspective, this means that functions cannot have side effects except for printing and changing the weight (probability) of a simulation. Also, the code binds variable names to values, and these values cannot change when running the program (immutability). Introducing a new binding with the same name hides the old binding (shadowing). These features greatly facilitate the application of automatic inference algorithms. Most computational biologists should be somewhat familiar with the functional programming style, as it is supported in both R and Python, even though the enforcement of immutability may require some adjustment of coding practices.

#### Algebraic data types

TreePPL uses a flexible system for data types, allowing users to define their own composite types. For instance, consider the representation of phylogenetic trees, a type provided in the standard library:

**Listing 6:**
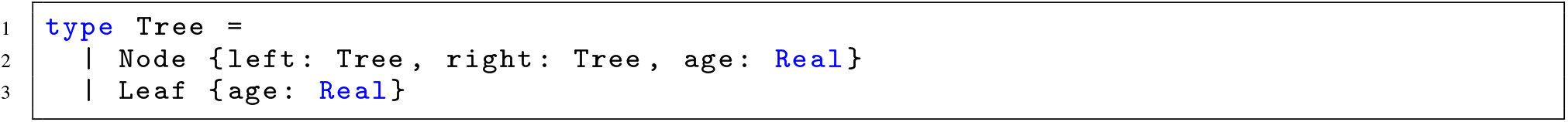
Definition of the `Tree` type in the standard library

The `Tree` type is a so-called *algebraic data type (ADT)* and is defined using two constructors: `Node` and `Leaf`. The Node constructor encapsulates a structure with two sub-trees (`left` and `right`) and an associated age. Conversely, the `Leaf` constructor represents terminal nodes with only an age attribute.

The specific type of a particular instance of an ADT can be checked using the `if … is` construct (see line 26 in the CRBD, Listing 4). This is a form of *pattern-matching* and checks if the instance is matched by a particular constructor. Pattern-matching allows different components of a phylogenetic tree to be handled in a natural way in TreePPL programs.

The basic Tree type can easily be extended by the user to cover more complex tree data structures with different model variables associated with its nodes and branches. New complex ADTs can be created by the user. The incorporation of ADTs in TreePPL provides a versatile framework for modeling intricate biological data structures, ensuring that the language remains both semantically rich and syntactically concise.

#### Type annotation

TreePPL is a statically typed language, which means that the type of each variable must be known at compile time; however, we only require the user to explicitly specify types in the function signatures using type annotation. Static typing using type annotation enables good error messages, readability, and certain compiler optimizations.

## 4 Scientific Modeling in TreePPL

In this section, we give three examples of modeling and inference in TreePPL. The examples represent very different types of phylogenetic models, and span a large portion of phylogenetic model space. They demonstrate a range of TreePPL modeling and inference features, and also highlight some challenges that remain for future work. The examples are by no means exhaustive. Users who want to describe their own models and use automated inference for them are encouraged to clone the TreePPL [GitHub repository](https://github.com/treeppl/treeppl/tree/main/models/phylo/clads) and explore the full set of examples provided there.

### 4.1 Diversification models

The CRBD model script used earlier in this paper forms a useful basis for exploring more complex diversification models, for which it is challenging to develop correct and efficient inference machinery. The basic techniques are the same as those recently used to analyze a range of diversification models accommodating different types of variation across lineages in diversification rates in the WebPPL and Birch PPLs [18]. In the TreePPL repository (in the `models/phylo/clads` directory), we provide an example of a more complex diversification model expressed as a TreePPL program, the cladogenetic diversification rate shift model (ClaDS) [58]. The difference between ClaDS and CRBD is that at each speciation (both observed and hidden), we propose a shift in the speciation and extinction rates in ClaDS. Otherwise, the general template for implementing it in TreePPL is the same as for the CRBD example discussed above.

The recent application of PPLs to diversification models made SMC inference algorithms available for these models for the first time [18]. Because SMC generates an estimate of the model likelihood as a byproduct, this opened up the possibility of resolving existing debates concerning the performance of different models in explaining empirical data using Bayesian model comparison. The TreePPL framework makes it possible to easily generate SMC inference machinery for a number of interesting diversification models that have been considered recently [59, 60], providing an effective way of assessing how well they explain reconstructed phylogenetic trees across different organism groups. For an up-to-date list of TreePPL diversification model examples, check [the on-line reference on phylogenetic models](https://treeppl.org/docs/Reference/phylo)).

### 4.2 Host repertoire model

The host repertoire model is arguably one of the most realistic models considered to date for the evolution of host-symbiont associations [61]. Given a set of host taxa, the host repertoire of a symbiont taxon is represented by an integer vector, where each element describes the interaction of the symbiont with each host taxon. For instance, one may consider three levels of interaction—non-host, potential host and actual host—as originally proposed when the model was introduced [61]. One can also consider a simpler model with just two levels of interaction—non-host and host—or more complex scenarios with more than three levels of interaction. Given a time-calibrated symbiont tree and a matrix of observed symbiont-host interactions as data, the model describes how the host repertoire evolves along the symbiont tree through changes in the level of interaction with each of the host taxa included in the interaction matrix.

Because ancestral host repertoires can include multiple hosts, this model can capture a wide range of complex scenarios of host-symbiont coevolution, from strict specialization in one-to-one interactions to complex pulses between generalist and specialist phases in the evolution of host ranges. Finally, with this model, one can also test if the probability of gaining a new host depends on the phylogenetic distance between the new host and the host(s) used at the time of the event. Originally proposed to study the evolution of butterfly-plant associations, this model is now being used for a variety of host-symbiont systems, for instance in analyses of beetle microbiomes [62] and parasitic mussels in fishes [63], as well as for other types of ecological interaction such as mimicry [64, 65].

The host repertoire model poses several interesting modeling and inference challenges. First, the number of possible host repertoires is huge. For a model with *n* hosts and three different levels of host interaction, there are 3^*n*^ *™*2^*n*^ possible host repertoires. The rate of change from one repertoire to another is computed from the loss and gain rates for the different levels of host interaction, potentially taking into account the phylogenetic dissimilarity between hosts (Figure 4). The size of the rate matrix means that we cannot integrate out ancestral host repertoires and changes between them for realistic-size problems, like we would do, say, for a standard substitution model describing the evolution of nucleotides in a DNA sequence. Instead, we need to sample over ancestral repertoires and histories of change between them.

**Figure 4:**
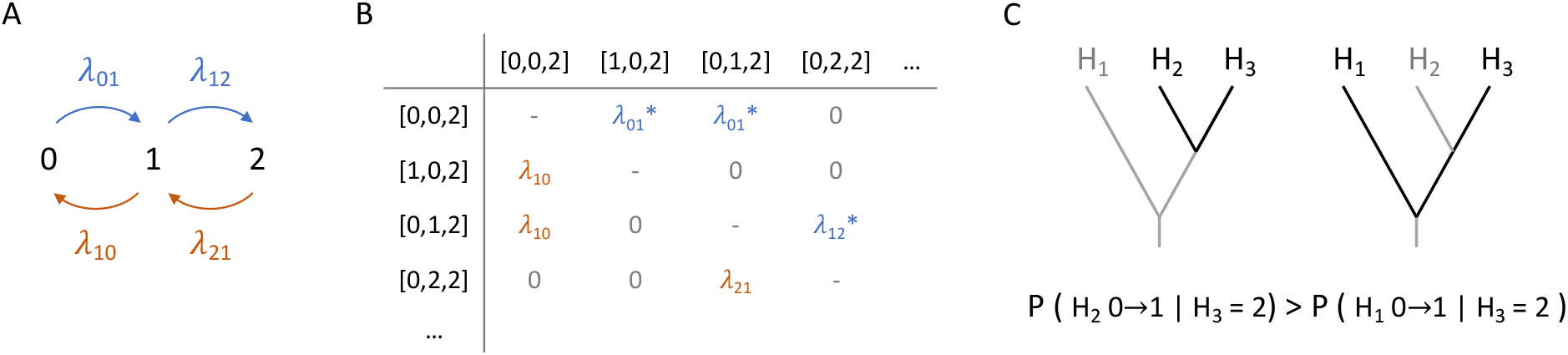
A. In the host repertoire model, different loss and gain rates are used to describe the rates of moving between levels of host interaction. B. A section of the rate matrix for a repertoire with three hosts, where all possible events have one of the four rates shown in A. Asterisks mark the gain rates, which are modified by a function of the phylogenetic distance between hosts. C. Phylogenetic distances between hosts. Since H2 is closer to H3 (the only host being used), the probability of gaining H2 is higher than gaining H1.

Second, it is challenging to sample ancestral host repertoires and change histories directly from the full model. It is more natural to sample from a simpler model, the independence model, in which host interactions evolve independently from each other. This is the same as sampling from *n* independently evolving discrete characters, each representing a different host. Each character would have three possible states corresponding to the different levels of host interaction. We can sample from this model, conditioned on the tip states, using well-known algorithms [66]. This is done by drawing from the appropriate probability distributions using *assume* statements (Figure 5). We then weight the sampled history according to its probability under the full model, factoring out the probability of the proposed state under the independence model. This is accomplished using *weight* statements, as illustrated in line 21 of Figure 5A.

**Figure 5:**
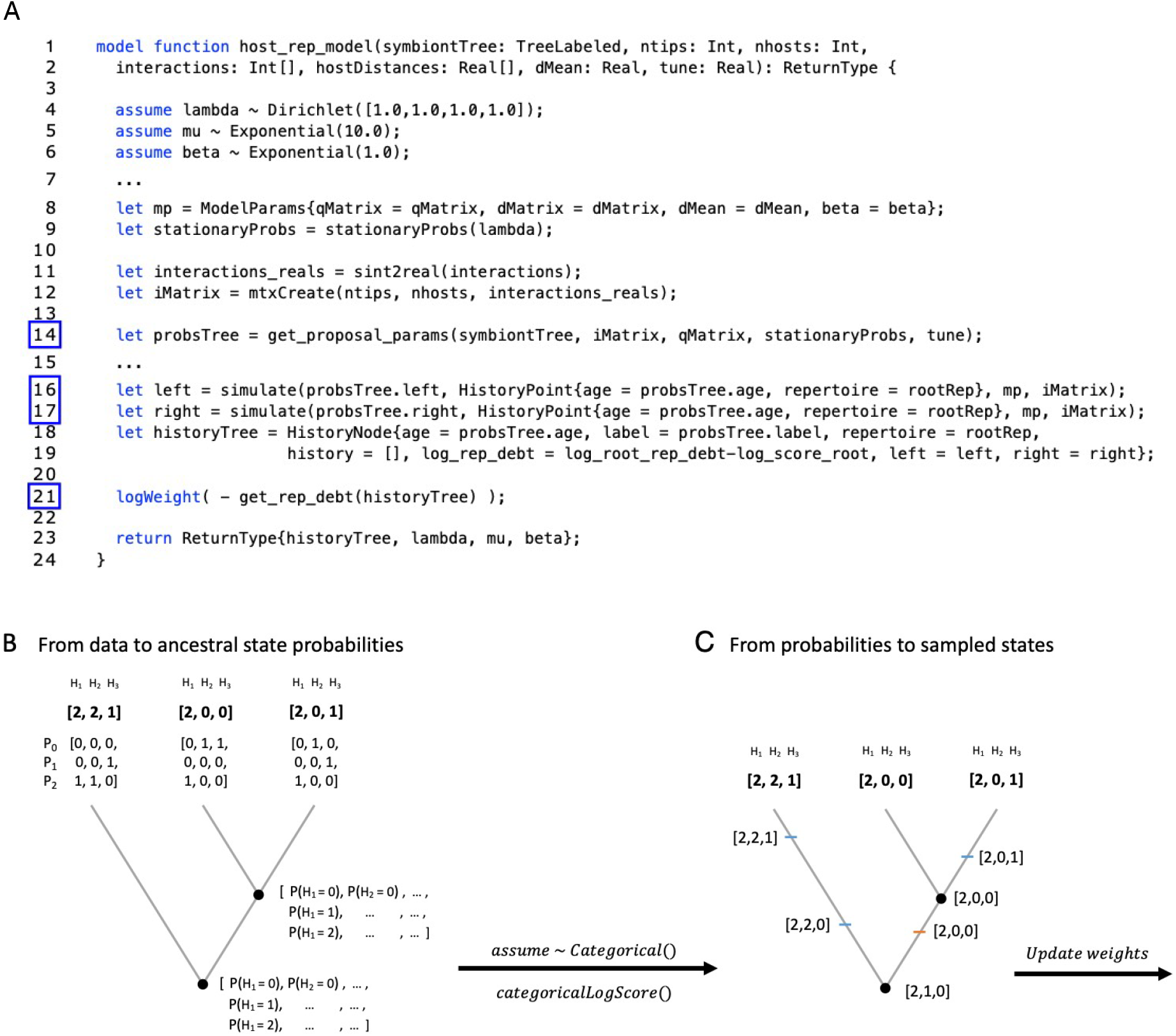
A. Simplified TreePPL program for inference of host repertoire evolution. In order to sample ancestral host repertoires, first (B) ancestral state probabilities are calculated based on the observed data and the independence model (line 14 in A). Then (C) a sample is drawn using *assume* statements and the probability weight of the sample is recorded (lines 16-17 in A). Lastly, probability weights are updated by factoring in the probability under the full model (also in lines 16-17 in A) and factoring out the recorded probability under the independence model (line 21 in A).

Because of the complexity of the host repertoire model, it is quite challenging to find efficient inference strategies. The only current implementation uses dedicated Markov chain Monte Carlo (MCMC) algorithms coded in C++ in the back end of the RevBayes platform [61]. Unfortunately, the host repertoire model is hard for the MCMC algorithm because of the difficulties of moving from one plausible history of repertoire change to another. Once the model is described in TreePPL, however, we get access to the full range of TreePPL inference strategies. They include sequential Monte Carlo (SMC) algorithms, which do not depend on moving between points in model space. TreePPL also supports mixed strategies involving both SMC and MCMC steps, and opens up for future inference strategies developed for all TreePPL models.

By modifying the TreePPL description of the model, we can easily extend the original model. For instance, one can associate an observation of the host repertoire of a leaf in the symbiont tree with some uncertainty about the true level of interaction with each host, make different assumptions about the root state (for example, assuming stationarity or not), or change the number of host interaction levels in the model.

### 4.3 Tree inference

To describe the problem of tree inference in a PPL, such that the code lends itself to automated inference using generic methods, we use stochastic recursion. The core idea is to control a recursive function using a random variable, such that successive iterations generate a valid draw from the prior probability distribution over tree space.

The PPL code accomplishing this can be structured as a forward simulation, in which the tree is generated from the root and forward in time until it is complete with all the observed leaves. It can also be structured as a backward simulation, in which one starts with all the leaves and simulates coalescent events backward in time until the root is reached. The latter, the coalescent process, is a more convenient description for our purposes, because it can be used to condition a simulation on observed data earlier than would otherwise be possible. For instance, in an SMC algorithm, the backward simulation makes it possible to weight a particle describing a subtree according to the probability of the substitution process generating the tip sequences on that subtree. In an MCMC algorithm, conditioning on the observed sequences early on in the simulation increases the probability of generating a complete tree that represents a good starting point for the Markov chain. Crucially for the end user, the model remains the same regardless of which inference strategy is used.

The disadvantage of simulating the process backwards is that we need to say something about ancestral DNA sequences in the tree before we can fully express the probability distribution they are drawn from. We solve this problem by using a proposal distribution, similar to the mechanism used to simulate from the host repertoire model using an initial draw from the independence model.

Assume, for simplicity, that we are modeling the evolution of DNA sequences using the Jukes-Cantor (JC) model. The coalescent process would start with a set of leaves, each leaf being associated with a DNA sequence. The leaves together form a set of trees, also known as a forest. This is expressed in TreePPL as a sequence of `Tree` variables, each `Tree` initially being an instance of a `Leaf` variable. A random pair is then drawn from this tree set, see line 6 of the code in Figure 6, and a coalescent time is drawn from the prior, see line 10.

**Figure 6:**
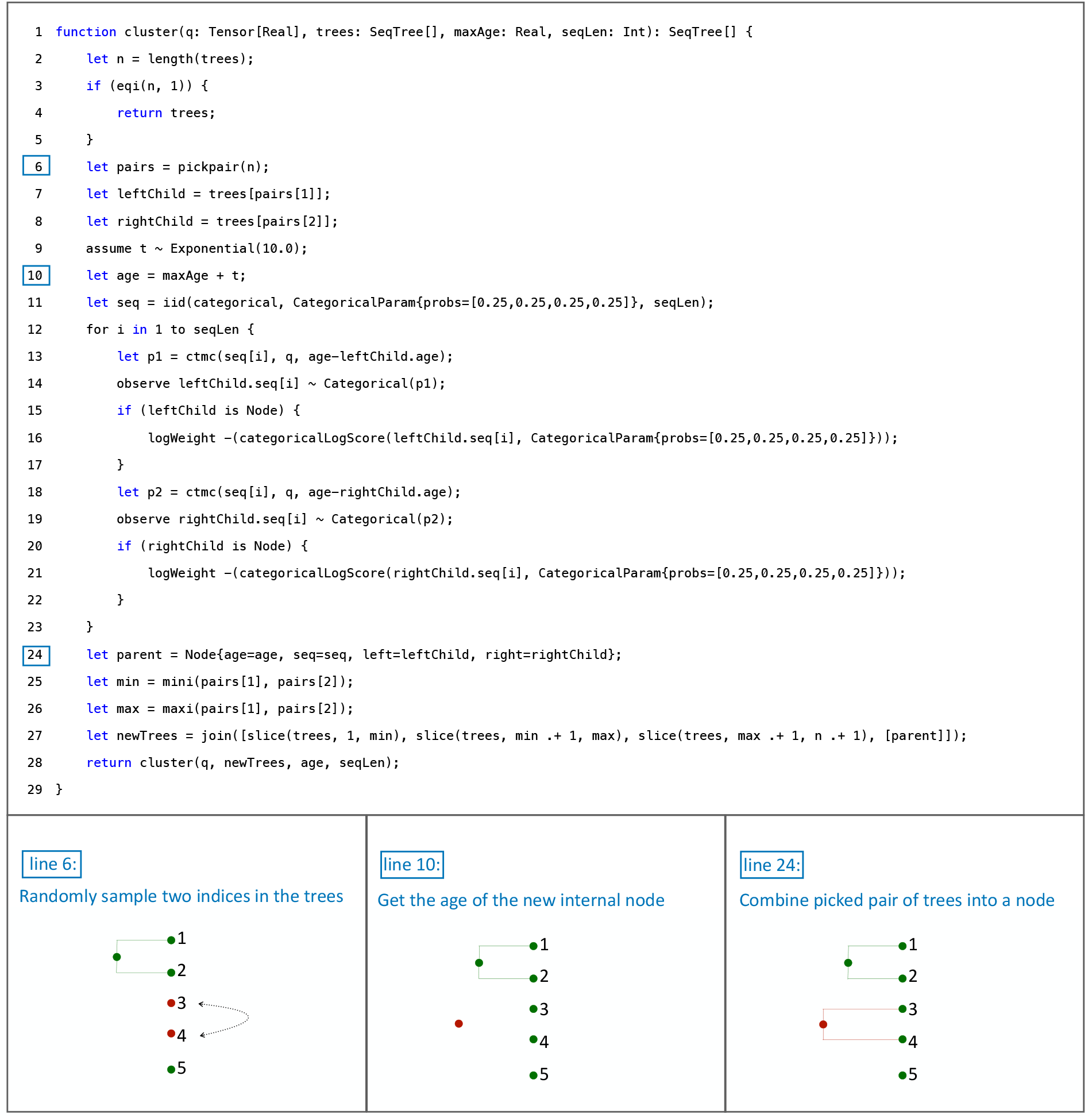
Illustration of the cluster function, used in phylogenetic tree inference.

Now we propose an ancestral DNA sequence from the stationary state distribution of the JC model, that is, we draw each nucleotide from a uniform distribution over the state set {*A, C, G, T*,} see line 11. Even though this is not the correct distribution, we correct the weight of the simulation later, when we know the correct probability distribution, just as in the host repertoire model. In principle, we could use any distribution for this proposal, as long as the weight of the simulation is later adjusted accordingly.

Once the ancestral DNA sequence has been specified, we can describe the probability of the two descendant sequences using the appropriate distribution from the continuous time Markov chain defined by the JC model over the appropriate time interval (lines 12–23 in Figure 6). If the descendant is a leaf, this is done using an observe statement. If it is an internal node, however, its DNA sequence has been proposed in an earlier pass through the function as a draw from the stationary distribution of the JC model. Therefore, we correct the weight of the simulation by factoring out the original draw and factoring in the probability of drawing the sequence from the correct continuous-time Markov chain distribution (see lines 16 and 21).

The program described above defines the correct probability distribution but current automated inference algorithms do not scale very well when applied to this program because they will sample over all possible ancestral DNA sequences. This is inefficient, as the sampled parameter space will be huge even for DNA sequences of moderate length. The standard solution is to compute tree probabilities by summing over all possible ancestral sequences using Felsenstein’s pruning algorithm [67], effectively removing the ancestral sequences from the sampled space. In TreePPL, the pruning algorithm can be expressed explicitly, and for that model the current version of TreePPL supports efficient SMC inference of phylogeny as described by [33]. In ongoing work [57], we are exploring the possibility of automatically implementing the pruning algorithm as part of the inference machinery, so that the user does not have to worry about expressing it in the model script.

The tree inference script described above, with the pruning algorithm, can be modified to support a range of related inference problems, such as inference of substitution model parameters, phylogenetic placement of unknown sequences in a known phylogenetic tree, or online inference of phylogeny, in which sequences are added sequentially to a growing phylogenetic tree. We point the interested reader to the TreePPL website for an up-to-date set of such model scripts. A pair of core tree inference models in TreePPL was verified for correctness against MrBayes (see Supplementary Figure S1).

Finally, we note that the current version of TreePPL also supports MCMC inference of tree topology from the same script. Again, with suitable proposal distributions, the inference should be efficient. However, the current MCMC algorithm does not support the stochastic tree topology moves that are currently used in most Bayesian MCMC phylogenetic software, such as stochastic nearest neighbor interchange or subtree pruning and regrafting [68]. Extending the automatic inference to support these techniques is left for future work.

## 5 Discussion and Conclusions

Clearly, the PPL approach comes with numerous benefits for the end user. Most importantly, perhaps, the clean separation of modeling from inference allows a domain expert to rapidly explore alternative models that could explain a phenomenon of interest, rather than spending considerable time and resources on implementing inference machinery from scratch for each and every model explored. This aspect is particularly important when scientific progress is tied to the exploration of new models that have not been considered previously, as is often the case in phylogenetics and evolutionary biology. Another important advantage of separating inference from modeling is that it makes it possible to reuse inference algorithms that have been extensively used and tested on other scientific modeling and inference problems, thereby reducing the risk of inference errors. Our examples provide some illustration of how a universal PPL, such as TreePPL, can be used to describe a range of common inference problems in phylogenetics in a succinct and intuitive way.

The potential disadvantage of the PPL approach is that the available automatic inference algorithms may be less efficient than dedicated inference algorithms written for a specific problem. Sometimes the difference is small enough that it makes no difference in practice. In other cases, the convenience of automatic inference and the speed with which it allows new models to be explored and applied to real data outweighs the loss in inference efficiency. Nevertheless, it is also true that there are many problem instances where the current automatic inference algorithms are simply too slow to allow realistic-size problems to be addressed. Research into better algorithms is currently very active, and is successively addressing these problems. Importantly, such advances are applicable across scientific domains, making it possible for PPL users to benefit from them as soon as they become available in the PPL they are using.

Meanwhile, as demonstrated in this paper and elsewhere [18], it is often possible to structure a PPL description so that it lends itself to efficient inference with current automatic inference techniques, before the appropriate solutions that would allow a more natural model description become available in PPL compilers. In other cases, it is possible to manually provide hints to the compiler, for instance on suitable resampling points, improving the efficiency of the generated inference machinery. These are not ideal solutions, but they vastly expand the utility of current PPLs, and provide excellent guidance as to relevant directions in future work on automatic inference techniques.

It is usually the conditioning on observed data, which makes the application of automatic inference techniques to probabilistic programs difficult. With some understanding of how automatic inference techniques work, it is possible to avoid many problems with inefficient inference in PPLs by structuring the model code appropriately. For instance, it is often advantageous for automatic inference to condition on observed data early on in the simulation, if possible. The host repertoire program accomplishes this by using the observed data already in deriving a proposal distribution that is similar to the posterior probability distribution. The tree inference program instead reverses the time flow in the simulation, so that it starts with the observed data and goes backwards in time.

We have designed TreePPL to support evolutionary biologists interested in exploring PPLs for phylogenetic problem instances. Primary goals have been to make it easy to describe phylogenetic models, and to provide an inference toolbox that includes state-of-the-art techniques providing efficient inference for such models. Some TreePPL language features may appear exotic to users in the biology community, such as algebraic data types and the adoption of an immutable functional programming style. These features are used to enable efficient inference, but they also introduce abstractions and programming practices that are powerful in expressing phylogenetic models, and therefore worth the learning effort. The advanced features are typically inspired by typed functional languages, such as OCaml, and provide a natural bridge to the CorePPL language used to express automatic inference algorithms in the Miking CorePPL framework. Importantly, the TreePPL language comes with a set of built-in types and functions, which facilitate the specification of phylogenetic models. For instance, the algebraic data type is easy to use in describing simple phylogenetic trees, and in extending these descriptions for models using more complex tree structures with many latent branch and node variables. The functions include operations that are commonly used in phylogenetics, such as counting the number of leaves on a tree, computing total tree length, or rotating subtrees. Finally, the TreePPL web site provides a number of example programs describing common phylogenetic simulation and inference problems beyond the ones discussed in this paper.

The TreePPL Python package offers powerful utilities for pre-processing and post-processing, and for controlling and running TreePPL programs. This opens up the possibility of using a wide range of Python tools developed specifically for data processing and visualization of data and results relevant for the phylogenetics community. The R package for TreePPL provides similar functionality for R users.

With respect to automatic inference algorithms, the current version of TreePPL implements a number of techniques, which have been found to be important for phylogenetic problems but that are not yet widely available in PPLs. These include selective CPS transformation [19], automatic alignment [30] and suspension analysis [20], and the alive particle filter [31]. Further examples of important techniques for efficient inference on phylogenetic problems include automatic application of delayed sampling, Felsenstein’s pruning algorithm, and effective MCMC samplers of tree topology. These techniques are currently all work in progress in TreePPL. In addition to these specialized techniques, TreePPL includes a range of more commonly supported automatic inference techniques for PPLs, some of which have rarely been used for phylogenetic problems previously.

There is currently a large set of generic PPLs, including Stan [69], Gen [70, 71], Edward [72], Turing [73], Anglican [22], WebPPL [21], Birch [74] and Pyro [75], to name a few. Arguably, none of them supports phylogenetic model specification as well as TreePPL, and—to our knowledge—none of them provides such a complete range of tools for efficient automatic inference on phylogenetic models. In addition, the TreePPL interface to Python and R facilitates the preprocessing of data and postprocessing of inference results for evolutionary biologists used to these environments, and all the support they provide for handling phylogenetics-related tasks.

Nevertheless, several of the more generic PPLs have large and active user communities, and their capabilities expand very rapidly. Several of them have been shown to be useful for phylogenetic problems already with their current set of features [18, 11]. Therefore, we strongly recommend phylogeneticists and evolutionary biologists interested in the PPL approach to also keep an eye on these more generic PPLs. Eventually, we will probably see the different PPL sub-communities converge on a smaller set of languages and inference compilers, but there are likely to be several years of intense parallel development of PPL techniques before the field matures in this way.

We would like to end with a discussion of the future of PPLs. At the core of probabilistic programming lies the Bayesian approach, which unites learning from observational data (supplied by the experiment) with prior belief (an aggregate belief from previous experiments) and a model (formulated by the scientist). When it comes to inference, the Bayesian approach is inherently probabilistic and assigns a (posterior) probability to the predictions that the model makes. This can be contrasted to neural networks that learn a high-dimensional model strictly from the observational data by optimizing a gradient function and are not based on explicit modeling of real-world phenomena.

We argue that a probabilistic approach is well-suited for scientific purposes as the data are not always conclusive, and we often need to weigh several competing hypotheses against each other. More importantly, a PPL model is based on a mechanistic understanding of the involved phenomena, as it can be used not only for inference but also generatively to simulate data under the model. The mechanistic model provides a succinct and accessible representation of the real world. Neural networks often lack interpretability and do not provide insights into the underlying mechanisms, which is crucial in scientific research. In contrast, a model described as a probabilistic program can be studied with analytical tools, and theorems can be proven about it—–we are not limited to only statistically analyzing its output.

In this sense, PPL approaches could be used in science in areas where there is insufficient understanding of the mechanisms of a phenomenon and where data are incomplete or noisy and can support several alternative explanations, such as is the case in evolutionary biology. For instance, many phylogenomic data sets come with poorly characterized systematic errors that originated during the sequencing and genome assembly steps. These errors can cause serious problems in phylogenomic inference, as they may give rise to a strongly misleading signal when probabilistic inference under evolutionary models is applied to big data sets [76, 77].

Machine learning could be used to improve the inference performance of PPLs. For instance, auto-tuning the inference algorithms is a problem that appears well suited to machine learning, with the caveat that the conditions for algorithm correctness must be respected. Finding an optimal inference strategy by selecting among a predefined set of techniques or possible combinations of them is a similar but more challenging problem. For early work in this direction for maximum likelihood optimization algorithms, see [78].

In summary, we contend that PPLs will have a significant impact on the future development and use of the next generation of phylogenetic models and inference problems. TreePPL is an attempt to create a simple and powerful language with an efficient software implementation that makes it easier and faster for biologists to solve relevant tasks. Yet, many interesting challenges remain to be solved, particularly related to scalability and performance, where techniques that combine advancements in probabilistic inference, machine learning, compiler optimizations, and phylogenetic domain knowledge are vital for success.

## Acknowledgements

This project was in part financially supported by the Swedish Foundation for Strategic Research (FFL15–0032 and RIT15–0012), by Digital Futures and by the Swedish Research Council (grant 2018–04329 awarded to DB, grants 2018-04620 and 2021–04830 awarded to FR, and International Postdoc Grant 2020–06422 awarded to MPB), and by the European Union’s Horizon 2020 research and innovation program under the Marie Skłodowska–Curie grant agreement PhyPPL No. 898120 to V.S. The research has also partially been carried out as part of the Vinnova Competence Center for Trustworthy Edge Computing Systems and Applications at KTH Royal Institute of Technology. This work was also partially supported by the Wallenberg AI, Autonomous Systems and Software Program (WASP) funded by the Knut and Alice Wallenberg Foundation.

## TreePPL Technical Details

| Item | Details |
| --- | --- |
| Version Number | TreePPL 2024 release candidate 1 |
| Webpage | https://treeppl.org |
| Download Site – Language | https://github.com/treeppl/treeppl |
| Download Site – Python environment | https://github.com/treeppl/treeppl-python |
| Download Site – R package | https://github.com/treeppl/treepplr |

## Footnotes

3 https://github.com/treeppl/treeppl-python

4 https://github.com/treeppl/treepplr

5 https://github.com/kudlicka/nexus2phyjson/blob/master/doc/phyjson_format_description.md

