## Supplementary Material for "TreePPL: A Universal Probabilistic Programming Language for Phylogenetics"


**Viktor Senderov***
Institut de Biologie de l'École normale supérieure - PSL

**Jan Kudlicka***
Department of Data Science and Analytics, BI Norwegian Business School

**Daniel Lundén**
Oracle

**Viktor Palmkvist**
EECS and Digital Futures, KTH Royal Institute of Technology

**Mariana P. Braga**
Department of Ecology, Swedish University of Agricultural Sciences

**Emma Granqvist**
Department of Bioinformatics and Genetics, Swedish Museum of Natural History

**Gizem Çaylak**
EECS and Digital Futures, KTH Royal Institute of Technology

**Thimothée Virgoulay**
Department of Bioinformatics and Genetics, Swedish Museum of Natural History

**David Broman†**
EECS and Digital Futures, KTH Royal Institute of Technology

**Fredrik Ronquist†**
Department of Bioinformatics and Genetics, Swedish Museum of Natural History
``



November 13, 2024


---

*Equal first author contributions between VS and JK.
†Equal senior author contributions between DB and FR.

### Contents

**1 Verification of tree inference models** **2**

**2 Verification of host repertoire model** **3**

#### 1 Verification of tree inference models

The strategy of tree inference in TreePPL using SMC was introduced in section 3.3. To verify the correctness of the TreePPL implementation of tree inference models, a set of core models have been verified against the commonly used MrBayes software [1].

These core TreePPL models use Felsenstein's pruning algorithm [2], and the evolution of the DNA sequences is either modelled by the Jukes-Cantor (JC) model or the General Time Reversible (GTR) model. See [3] for a comprehensive overview of nucleotide substitution models. A small test dataset was designed for testing purposes, with 4 leaves and 15 sites.

The marginal likelihood under the different models was computed with SMC in TreePPL. The same dataset, and the same assumptions in terms of nucleotide substitution models and priors, was used in MrBayes. The inference method in MrBayes is MCMC, and the standard output delivers the harmonic mean which can be a good approximation for the marginal likelihood. However, for a more exact approximation MrBayes also provides the stepping-stone algorithm [4]. In Figure S1 below, both the harmonic mean and the marginal likelihood from MrBayes is shown alongside the marginal likelihood from TreePPL. Figure S1A shows results under the JC model, figure S1B show results under the GTR model.

The results verify that the implemented TreePPL models deliver the same marginal likelihood as MrBayes, and with a higher precision. The Harmonic mean estimates are higher in both cases, which is expected. See for example MrBayes version 3.2 Manual section 4.5 for further details on the difference between harmonic mean and stepping-stone estimates.

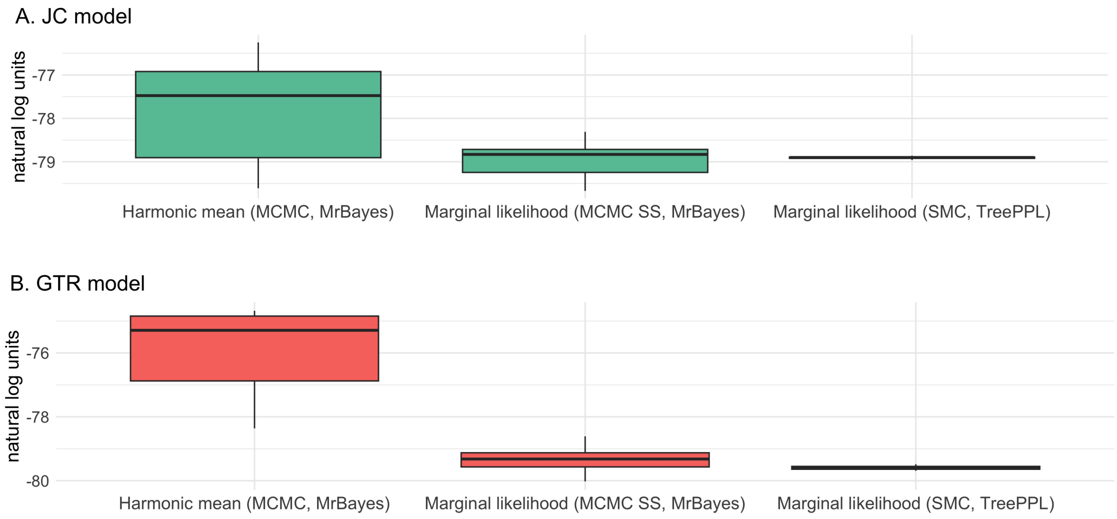


Figure S1: Comparing results between TreePPL and MrBayes: the harmonic mean or the marginal likelihood depending on the inference method used. Figure S1A shows the results under the Jukes-Cantor (JC) model, and S1B under the general time reversible (GTR) model. The boxplots contain the result of ten runs for each method respectively.

#### 2 Verification of host repertoire model

To verify the implementation of the host repertoire model in TreePPL, we ran the same analysis in TreePPL and RevBayes (the software where the model was first implemented) and compared posterior distributions. We created a small dataset with actual and potential interactions between 3 symbionts and 3 hosts (Figure S2). The interaction matrix was composed of 5 actual interactions and 1 potential interaction. We then estimated the joint posterior distribution of model parameters and evolutionary histories using the inference strategy described in [5].

In RevBayes, we ran 2 independent MCMC analyses, for $10^6$ cycles, sampling parameters and character histories every 100 cycles, and discarding the first 10% as burnin. Results from a single MCMC analysis are presented given that convergence was confirmed.

In TreePPL, we ran 1000 sweeps and took a subsample of 10 from a total of $3 \times 10^4$ particles from each sweep. That was the number of particles necessary to achieve convergence between sweeps. We used variance of log normalizing constants close to 1 as the convergence criterion. The subsampling was done with replacement and probability proportional to the weight of the particle. Parameter densities were calculated based on the $10^4$ particles (1000 sweeps x 10), using the normalizing constant of each sweep as weight.

Inferred ancestral states were identical in TreePPL and RevBayes (Fig. S2), with differences in posterior probabilities of particular interactions too small to affect the results. There was more uncertainty in parameter estimation (Fig. S3), most likely because the dataset is small. The main difference between implementations was in the marginal posterior densities of the clock parameter. This difference is most likely due to transformations of the Q matrix done in RevBayes but not in TreePPL.

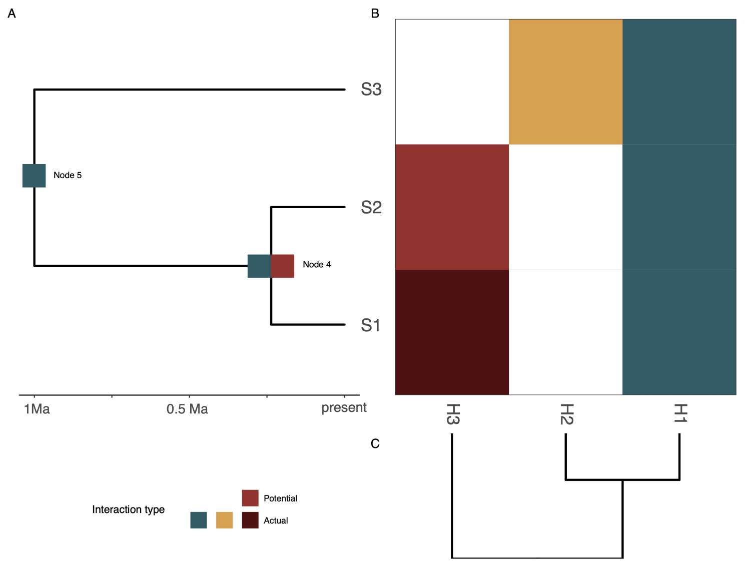


Figure S2: Simulated data and ancestral state reconstruction showing interactions with marginal posterior probability $\geq$ 0.9. The same result was produced by RevBayes and TreePPL. The model reconstructs how host repertoire evolved along the symbiont phylogeny (A), based on the observed (in this case, simulated) interactions (B), and the phylogenetic distance between hosts (C). The matrix in B shows the interactions between symbionts (rows) and hosts (columns) and cells are colored by host and by interaction type (the single potential interaction in the simulated data has a lighter shade than the actual interaction with host H3). Rows and columns are ordered to match the phylogenetic trees. Each square at the internal nodes of the symbiont tree represents an inferred ancestral interaction with one host.

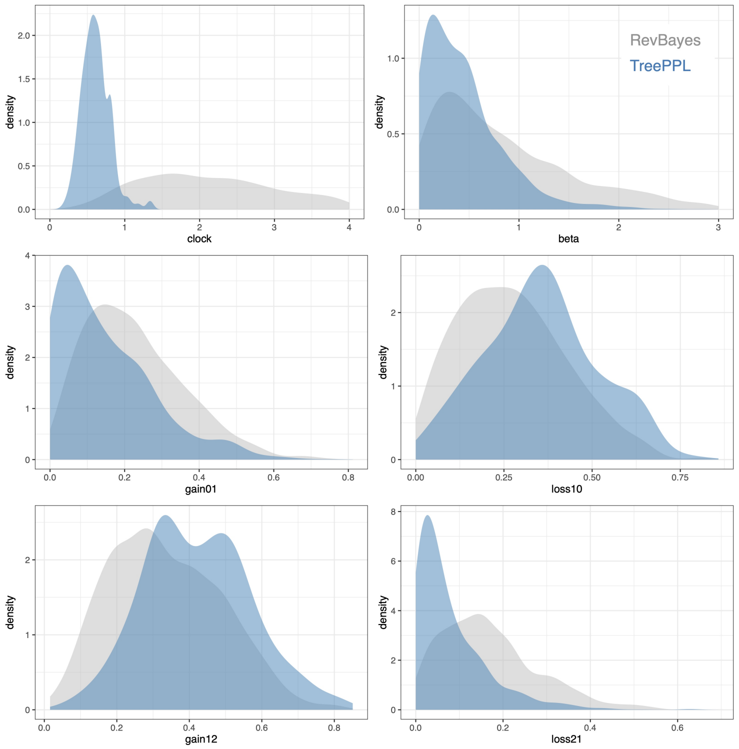


Figure S3: Marginal posterior densities for parameters of the host-repertoire model from the analysis of simulated data done in RevBayes (grey) and in TreePPL (blue).

[3] Ziheng Yang. *Molecular Evolution: A Statistical Approach*. Oxford University Press, 05 2014.

[4] Fredrik Ronquist, Maxim Teslenko, Paul Van Der Mark, Daniel L Ayres, Aaron Darling, Sebastian Höhna, Bret Larget, Liang Liu, Marc A Suchard, and John P Huelsenbeck. Mrbayes 3.2: efficient bayesian phylogenetic inference and model choice across a large model space. *Systematic biology*, 61(3):539–542, 2012.

[5] Mariana P Braga, Michael J Landis, Sören Nylin, Niklas Janz, and Fredrik Ronquist. Bayesian Inference of Ancestral Host-parasite Interactions under a Phylogenetic Model of Host Repertoire Evolution. *Systematic Biology*, 69(6):1149–1162, 2020.
